# Neutralization of SARS-CoV-2 Omicron sublineages by 4 doses of mRNA vaccine

**DOI:** 10.1101/2022.07.29.502055

**Authors:** Xuping Xie, Jing Zou, Chaitanya Kurhade, Mingru Liu, Ping Ren, Pei-Yong Shi

**Affiliations:** Department of Biochemistry and Molecular Biology, University of Texas Medical Branch, Galveston, TX, USA; Department of Pathology, University of Texas Medical Branch, Galveston, TX, USA; Institute for Human Infection and Immunity, University of Texas Medical Branch, Galveston, TX, USA; Sealy Institute for Drug Discovery, University of Texas Medical Branch, Galveston, TX, USA; Institute for Translational Sciences, University of Texas Medical Branch, Galveston, TX, USA; Sealy Institute for Vaccine Sciences, University of Texas Medical Branch, Galveston, TX, USA; Sealy Center for Structural Biology & Molecular Biophysics, University of Texas Medical Branch, Galveston, TX, USA

## Abstract

Since the initial emergence of SARS-CoV-2 Omicron BA.1, several Omicron sublineages have emerged, leading to BA.5 as the current dominant sublineage. Here we report the neutralization of different Omicron sublineages by human sera collected from individuals who had distinct mRNA vaccination and/or BA.1 infection. Four-dose-vaccine sera neutralize the original USA-WA1/2020, Omicron BA.1, BA.2, BA.212.1, BA.3, and BA.4/5 viruses with geometric mean titers (GMTs) of 1554, 357, 236, 236, 165, and 95, respectively; 2-dose-vaccine-plus-BA.1-infection sera exhibit GMTs of 2114, 1705, 730, 961, 813, and 274, respectively; and 3-dose-vaccine-plus-BA.1-infection sera show GMTs of 2962, 2038, 983, 1190, 1019, and 297, respectively. Thus, 4-dose-vaccine elicits the lowest neutralization against BA.5; 2-dose-vaccine-plus-BA.1-infection elicits significantly higher GMTs against Omicron sublineages than 4-dose-vaccine; and 3-dose-vaccine-plus-BA.1-infection elicits slightly higher GMTs (statistically insignificant) than the 2-dose-vaccine-plus-BA.1-infection. Finally, compared with BA.5, the newly emerged BA.2.75 is equally evasive of 4-dose-vaccine-elicited neutralization, but more susceptible to 3-dose-vaccine-plus-BA.1-infection-elicited neutralization.

## Introduction

The emergence of severe acute respiratory syndrome coronavirus 2 (SARS-CoV-2) Omicron variant was first identified in South Africa in November 2021. Due to Omicron’s improved viral transmission and immune evasion^1,2^, the World Health Organization designated it as the fifth variant of concern (VOC) after the previous Alpha, Beta, Gamma, and Delta VOCs. Since then, the Omicron variant has evolved to several sublineages, including BA.1, BA.2, BA.2.12.1, BA.3, BA.4, and BA.5. Among these sublineages, only Omicron BA.3 remained at a low frequency in circulation, most likely due to its low fitness; whereas other sublineages sequentially increased their prevalence over time. As of July 16, 2022 in the United States, sublineage BA.2.12.1, BA.4, and BA.5 accounted for 8.6%, 12.8%, and 77.9% of the total COVID-19 cases, respectively (https://covid.cdc.gov/covid-datatracker/#variant-proportions). Besides the above Omicron sublineages, a new sublineage BA.2.75 emerged in late May 2022 and has increased its prevalence in many countries. It is thus important to determine the neutralization susceptibility of the ongoing Omicron sublineages, particularly the most prevalent BA.5, to vaccination and previous infections.

Immediately after the emergence of the Omicron variant, the first identified sublineage BA.1 was found to evade vaccine-elicited neutralization more efficiently than any previous VOCs^2-7^. Two doses of mRNA vaccine did not elicit robust neutralization against BA.1; 3 doses of vaccine were required to generate sufficient neutralization against BA.1^8^. Non-Omicron infection did not elicit robust neutralization against BA.1 either^9^. Among all the Omicron sublineages, BA.5 exhibited the greatest evasion of vaccine-mediated neutralization; 3 doses of BNT162b2 mRNA vaccine elicited weak neutralization against BA.5^10^. The latter result, together with the high prevalence of BA.5, underscores the urgency to examine the neutralization of BA.5 after four doses of mRNA vaccine. In addition, many people who had previously received two or three doses of vaccine contracted BA.1 breakthrough infection during the initial Omicron wave; it would be important to evaluate their antibody neutralization against BA.5. To address these key questions, we have characterized the neutralization profiles of sera, obtained from humans with distinct mRNA vaccination and/or BA.1 infection, against different Omicron sublineages.

## Results and discussions

### Experimental approach and rationale

We used a set of previously established chimeric SARS-CoV-2s to determine the serum neutralization against different Omicron sublineages. Each chimeric SARS-CoV-2 contained a complete spike gene from BA.1, BA.2, BA.2.12.1, BA.3, or BA.4/5 in the backbone of USA-WA1/2020 (a virus strain isolated in January 2020), resulting in BA.1-, BA.2-, BA.2.12.1-, BA.3-, or BA.4/5-spike SARS-CoV-2^10^. BA.4 and BA.5 have an identical spike sequence and are denoted as BA.4/5. **Figure 1A** summarizes the amino acid mutations of the spike protein from different Omicron sublineages. An mNeonGreen (mNG) gene was engineered into the open-reading-frame-7 (ORF7) of the viral genome to enable a fluorescent focus reduction neutralization test (FFRNT) in a high-throughput format^9^. The FFRNT has been reliably used to measure antibody neutralization for COVID-19 vaccine research and development^6,10^.

**Figure 1.**
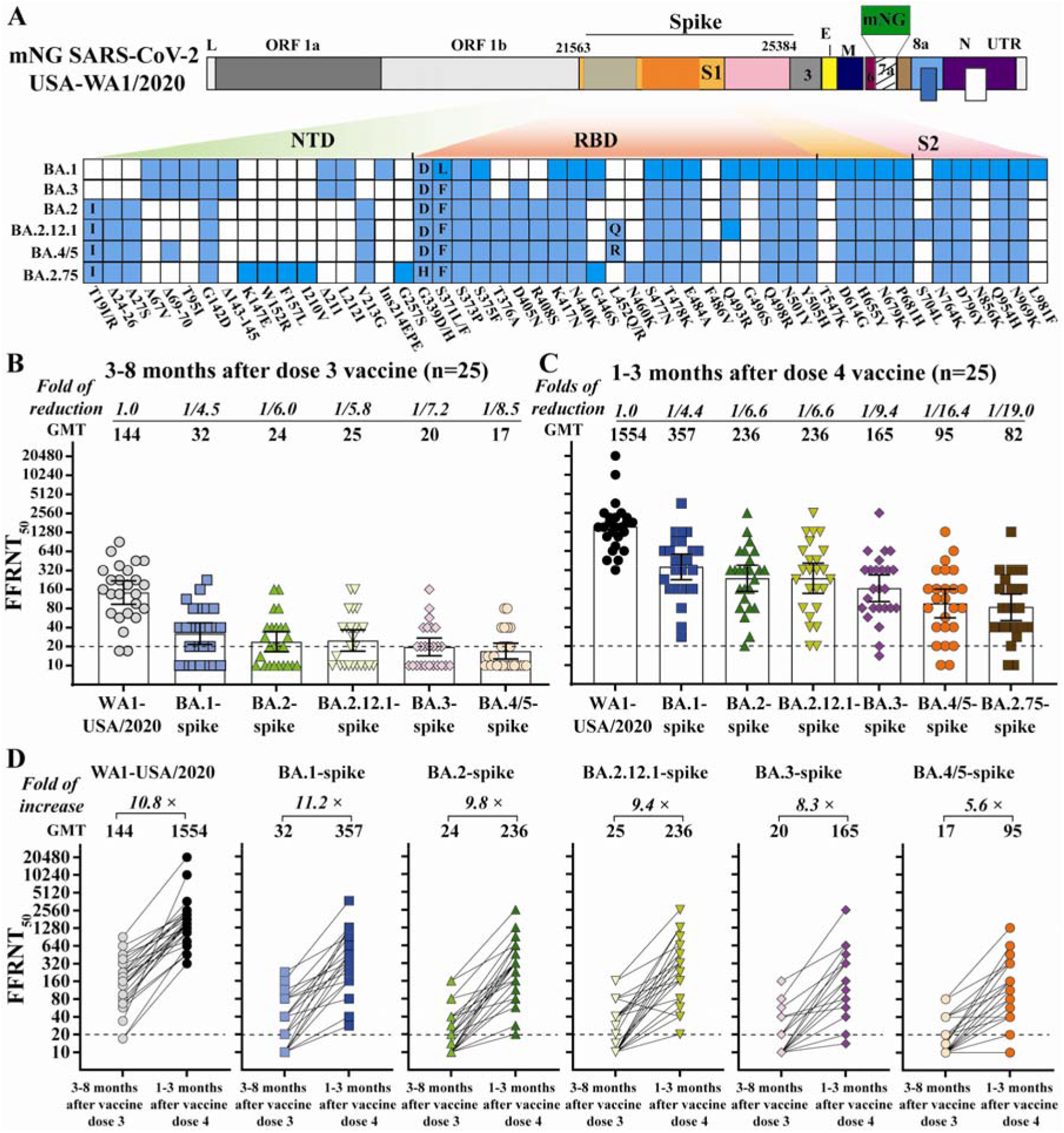
Neutralization of Omicron sublineages before and after 4 doses of mRNA vaccine. (**A**) Construction of Omicron sublineage-spike mNG SARS-CoV-2. mNG USA-WA1/2020 SARS-CoV-2 was used to engineer Omicron-spike SARS-CoV-2s. The mNG reporter gene was engineered at the open-reading-frame-7 (ORF7) of the USA-WA1/2020 genome.1 Amino acid mutations, deletions (Δ), and insertions (Ins) are indicated for variant spikes in reference to the USA-WA1/2020 spike. L: leader sequence; ORF: open reading frame; NTD: N-terminal domain of S1; RBD: receptor binding domain of S1; S: spike glycoprotein; S1: N-terminal furin cleavage fragment of S; S2: C-terminal furin cleavage fragment of S; E: envelope protein; M: membrane protein; N: nucleoprotein; UTR: untranslated region. Twenty-five pairs of human sera were collected 3-8 month after dose 3 and 1-3 month after dose 4 mRNA vaccine. The FFRNT_50_s for mNG BA.1-, BA.2-, BA.2.12.1, BA.3-, and BA.4/5-spike SARS-CoV-2s were determined in duplicate assays; the FFRNT_50_ for USA-WA1/2020 SARS-CoV-2 was determined in two independent experiments, each with duplicate assays. (**B**) FFRNT_50_ of sera collected before dose 4 vaccine. The bar heights and the numbers above indicate neutralizing GMTs. The whiskers indicate 95% CI. The fold of GMT reduction against each Omicron sublineage, compared with the GMT against USA-WA1/2020, is shown in italic font. The dotted line indicates the limit of detection of FFRNT_50_. The *p* values (Wilcoxon matched-pairs signed-rank test) for group comparison of GMTs are the following. USA-WA1/2020 versus all Omicron sublineage-spike: <0.0001; BA.1-spike versus BA.2-, BA.2.12.1-, BA.3-, BA.4/5-spike: 0.004, 0.0336, <0.0001, < 0.0001, respectively; BA.2-spike versus BA.2.12.1-, BA.3-, BA.4/5-spike: 0.5, 0.065, 0.0083, respectively. BA.2.12.1-spike versus BA.3-, BA.4/5-spike: 0.0098, 0.0002, respectively; BA.3-spike versus BA.4/5-spike: 0.156. (**C**) FFRNT_50_ of sera collected after dose 4 vaccine. The *p* values (Wilcoxon matched-pairs signed-rank test) for group comparison of GMTs are the following. USA-WA1/2020 versus all Omicron sublineage-spike: <0.0001; BA.1-spike versus BA.2-, BA.2.12.1-, BA.3-, BA.4/5, BA.2.75-spike: 0.008, 0.033, <0.0001, < 0.0001, < 0.0001, respectively; BA.2-spike versus BA.2.12.1-, BA.3-, BA.4/5, BA.2.75-spike: 0.12, <0.0001, < 0.0001, < 0.0001, respectively; BA.2.12.1-spike versus BA.3-, BA.4/5, BA.2.75-spike: 0.0002, <0.0001, <0.0001, respectively; BA.3-spike versus BA.4/5-, BA.2.75-spike: 0.0009, <0.0001, respectively; BA.4/5-spike versus BA.2.75-spike: 0.142. (**D**) FFRNT_50_ values with connected lines for each serum pair before and after dose 4 vaccine. The GMT fold increase before and after dose 4 is shown in italic font. The *p* values of GMT (Wilcoxon matched-pairs signed-rank test) before and after dose 4 vaccines are all <0.0001.

Using FFRNT, we measured the neutralization of three panels of human sera against the chimeric Omicron sublineage-spike SARS-CoV-2s. The first panel consisted of 25 pairs of sera collected from individuals before and after dose 4 of Pfizer or Moderna vaccine (**Table S1**). Those specimens were tested negative against viral nucleocapsid protein, suggesting those individuals had not been infected by SARS-CoV-2. The second and third serum panels were collected from individuals who had received 2 (n=29; **Table S2**) or 3 (n=38; **Table S3**) doses of mRNA vaccine and subsequently contracted Omicron BA.1 breakthrough infection. The BA.1 breakthrough infection was confirmed for each patient by sequencing viral RNA collected from nasopharyngeal swab samples. **Tables S1-3** summarize (i) the serum information and (ii) the 50% fluorescent focus-reduction neutralization titers (FFRNT_50_) against USA-WA1/2020, BA.1-, BA.2-, BA.2.12.1-, BA.3-, and BA.4/5-spike SARS-CoV-2s.

### Low neutralization of Omicron BA.5 after 4 doses of mRNA vaccine

To measure 4 doses of vaccine-elicited neutralization, we collected 25 pairs of sera from individuals before and after dose 4 of Pfizer or Moderna mRNA vaccine. For each serum pair, one sample was collected 3-8 months after dose 3 vaccine; the other sample was obtained from the same individual 1-3 months after dose 4 vaccine (**Table S1**). Before the 4^th^ dose vaccine, the 3-dose-vaccine sera neutralized USA-WA1/2020, BA.1-, BA.2-, BA.2.12.1-, BA.3-, and BA.4/5-spike mNG viruses with low geometric mean titers (GMTs) of 144, 32, 24, 25, 20, and 17, respectively (**Fig. 1B**); after the 4^th^ dose vaccine, the GMTs increased significantly to 1554, 357, 236, 236, 165, and 95, respectively (**Fig. 1C**); so, the 4^th^ dose vaccine increased the neutralization against the corresponding viruses by 10.8-, 11.2-, 9.8-, 9.4-, 8.3-, and 5.6-fold, respectively (**Fig. 1D**). Despite the neutralization increase after the 4^th^ dose vaccine, the GMTs against BA.1-, BA.2-, BA.2.12.1-, BA.3-, and BA.4/5-spike viruses were 4.4-, 6.6-, 6.6-, 9.4-, and 16.4-fold lower than the GMT against the USA-WA1/2020, respectively (**Fig. 1C**). These results support three conclusions. First, among the tested Omicron sublineages, BA.5 possesses the greatest evasion of vaccine-elicited neutralization. Second, the 4^th^ dose of mRNA vaccine does not elicit robust neutralization against BA.5, with a GMT of 95. A recent study reported a neutralizing titer of 70 as the threshold to prevent breakthrough infections of Delta variant^11^. Although the minimal neutralizing titer required to prevent BA.5 infection has not been determined, the low neutralization against BA.5 after dose 3 vaccine (GMT of 103 at 1-month post dose 3, reported by Kurhade et al.^10^) and dose 4 vaccine (GMT of 95 at 1- to 3-month post dose 4, reported here), together with the increased viral transmissibility, could account for the on-going surge of BA.5 around the world. Third, an updated vaccine that matches the highly immune-evasive and prevalent BA.5 spike is needed to mitigate the current and future Omicron surges. Our results support the U.S. Food and Drug Administration’s recommendation to include BA.5 spike for future COVID-19 vaccine booster doses.

### High neutralization against BA.5 and other Omicron sublineages after 2 or 3 doses of vaccine plus BA.1 infection

To compare with 4-dose-vaccine sera, we measured the neutralization against Omicron sublineages using sera collected from individuals who had received 2 or 3 doses of mRNA vaccine and subsequently contracted BA.1 infection (**Fig. 2**). **Tables S2** and **S3** summarize the FFRNT_50_ results for 2-dose-vaccine-plus-BA.1-infection sera and 3-dose-vaccine-plus-BA.1-infection sera, respectively. The 2-dose-vaccine-plus-BA.1-infection sera neutralized BA.1, BA.2, BA.212.1, BA.3, and BA.4/5 with GMTs of 2114, 1705, 730, 961, 813, and 274, respectively (**Fig. 2A**); the 3-dose-vaccine-plus-BA.1-infection sera showed slightly higher GMTs of 2962, 2038, 983, 1190, 1019, and 297, respectively (**Fig. 2B**). So, the GMT ratios between the 3-dose-vaccine-plus-BA.1-infection sera and 2-dose-vaccine-plus-BA.1-infection sera were 1.4, 1.2, 1.3, 1.2, 1.3, and 1.1 when neutralizing USA-WA1/2020, BA.1-, BA.2-, BA.212.1-, BA.3-, and BA.4/5-spike viruses, respectively; these GMT differences between the two serum groups were statistically insignificant, suggesting the extra dose of vaccine does not significantly boost neutralization for the 3-dose-vaccine-plus-BA.1-infection sera.

**Figure 2.**
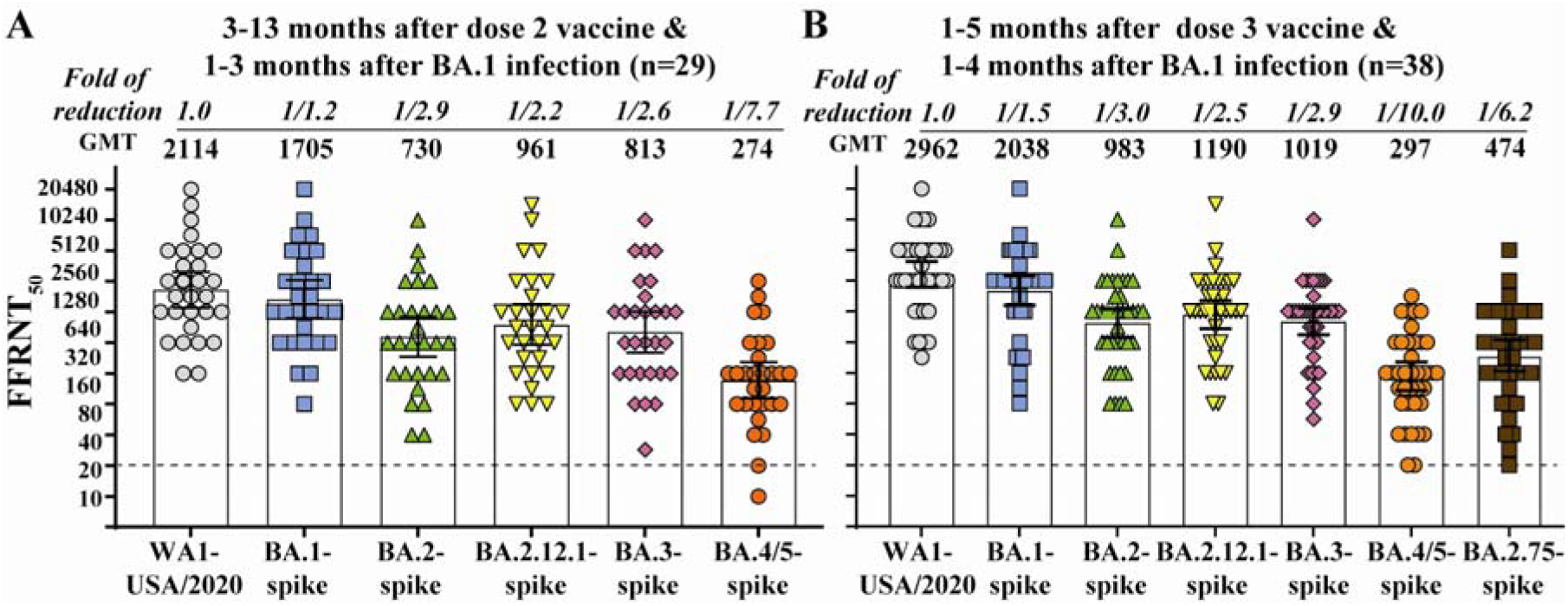
Neutralization of Omicron sublineages after 2 or 3 doses of mRNA vaccine and BA.1 infection. (**A**) FFRNT_50_ of 2-dose-vaccine-plus-BA.1-infection sera. Twenty-nine sera were collected from individuals who received 3 doses of vaccine and subsequently contracted BA.1 breakthrough infection. The GMT reduction fold against each Omicron sublineage and USA-WA1/2020 is shown in italic font. The dotted line indicates the limit of detection of FFRNT_50_. The *p* values (Wilcoxon matched-pairs signed-rank test) for group comparison of GMTs are indicated below. USA-WA1/2020 versus BA.1-spike, other sublineage-spikes: 0.053, <0.0001, respectively; BA.1-spike versus other sublineage-spike: <0.0001; BA.2-spike versus BA.2.12.1-, BA.3-, BA.4/5-spike: 0.0006, 0.0215, <0.0001, respectively; BA.2.12.1-spike versus BA.3- and BA.4/5-spike: 0.0309, <0.0001, respectively; BA.3-spike versus BA.4/5-spike SARS-CoV-2: <0.0001. (**B**) FFRNT_50_ of 2-dose-vaccine-plus-BA.1-infection sera. Thirty-eight sera were collected from individuals who received 3 doses of vaccine and subsequently contracted BA.1 infection. The *p* values (Wilcoxon matched-pairs signed-rank test) for group comparison of GMTs are indicated below. USA-WA1/2020 versus all Omicron sublineage-spike: <0.0001; BA.1-spike versus other sublineage-spike: <0.0001; BA.2-spike versus Omicron BA.2.12.1-, BA.3-, BA.4/5, BA.2.75-spike: 0.0082, 0.9095, <0.0001, <0.0001, respectively; BA.2.12.1-spike versus Omicron BA.3-, BA.4/5, BA.2.75-spike: 0.0018, <0.0001, <0.0001, respectively; BA.3-spike versus BA.4/5-, BA.2.75-spike: <0.0001, <0.0001, respectively; BA.4/5-spike versus BA.2.75-spike: 0.006.

In contrast, the GMT ratios between the 2-dose-vaccine-plus-BA.1-infection and 4-dose-vaccine sera were 1.4, 4.8, 3.1, 4.1, 4.9, and 3.9 when neutralizing USA-WA1/2020, BA.1-, BA.2-, BA.212.1-, BA.3-, and BA.4/5-spike viruses, respectively. The result suggests that, compared with the two extra doses of vaccine in the 4-dose-vaccine sera, the BA.1 infection in the 2-dose-vaccine-plus-BA.1-infection sera is more efficient in boosting both the magnitude and breadth of neutralization against all Omicron sublineages; however, the neutralization against BA.5 was still the lowest among all tested sublineages.

For the 2-dose-vaccine-plus-BA.1-infection sera, the GMTs against BA.1-, BA.2-, BA.2.12.1-, BA.3-, and BA.4/5-spike viruses were 1.2-, 2.9-, 2.2-, 2.6-, and 7.7-fold lower than the GMT against the USA-WA1/2020, respectively (**Fig. 2A**); similar results were observed for the 3-dose-vaccine-plus-BA.1-infection sera, with GMTs against BA.1-, BA.2-, BA.2.12.1-, BA.3-, and BA.4/5-spike viruses that were 1.5-, 3.0-, 2.5-, 2.9-, and 10-fold lower than the GMT against the USA-WA1/2020, respectively (**Fig. 2B**). The GMT decreases against Omicron sublineages for the 2-dose-vaccine-plus-BA.1-infection sera and those for the 3-dose-vaccine-plus-BA.1-infection sera are significantly less than those observed for the 4-dose-vaccine sera (Compare **Figs. 2A** and **2B** with **Fig. 1C**). The results again indicate that BA.1 infection of vaccinated people efficiently boosts the breadth of neutralization against all tested Omicron sublineages. However, such BA.1 infection-mediated boost of neutralizing magnitude/breadth is dependent on previous vaccination. This is because BA.1 infection of unvaccinated people did not elicit greater neutralizing magnitude/breadth against Omicron sublineages than 3 doses of mRNA vaccine^10^.

### Neutralization against Omicron sublineage BA.2.75

To assess the neutralization of the newly emerged Omicron sublineage BA.2.75, we engineered the complete spike gene of BA.2.75 (**Fig. 1A**) into the backbone of USA-WA1/2020, resulting in BA.2.75-spike SARS-CoV-2. The BA.2.75-spike SARS-CoV-2 was sequenced to ensure no undesired mutations. When tested with 4-dose-vaccine sera, the neutralizing GMTs against BA.2.75-spike virus (82) and BA.5-spike virus (95) were comparable (statistically insignificant; **Fig. 1C**), suggesting these two sublineages evade vaccine-elicited neutralization at a similar efficiency. However, when tested with 3-dose-vaccine-plus-BA.1-infection sera, the neutralizing GMT against BA.2.75-spike virus (474) was significantly higher than that against BA.5-spike virus (294; **Fig. 1C**), indicating that BA.1 infection boosts antibody neutralization differently from mRNA vaccination. These results further support the bivalent booster vaccine strategy.

### Study limitations

Our study has several limitations. First, this study lacks analysis of sera from vaccinated people who were infected with Omicron sublineages other than BA.1. It would be particularly important to study vaccinated individuals who contracted BA.5 infection. Second, we have not analyzed T cells and non-neutralizing antibodies that have Fc-mediated effector functions. These two immune arms, together with neutralizing antibodies, protect patients from severe disease. Many T cell epitopes after vaccination or natural infection are preserved in Omicron spikes^12^. Third, the relatively small sample size and heterogeneity in vaccination/infection time have weakened our ability to compare the results among the three distinct serum panels. Regardless of these limitations, our results consistently showed that (i) 4 doses of the current mRNA vaccine elicit low neutralization against BA.5 spike, rationalizing the need to update the vaccine sequence to match the highly immune-evasive and prevalent BA.5; (ii) vaccinated individuals with BA.1 break-through infection develop greater neutralizing magnitude/breadth against Omicron sublineages than those who have received 4 doses of mRNA vaccine. These laboratory investigations, together with real-world vaccine effectiveness data, will continue to guide vaccine strategy and public health policy.

## Methods

### Ethical statement

All virus work was performed in a biosafety level 3 (BSL-3) laboratory with redundant fans in the biosafety cabinets at The University of Texas Medical Branch at Galveston. All personnel wore powered air-purifying respirators (Breathe Easy, 3M) with Tyvek suits, aprons, booties, and double gloves.

The research protocol regarding the use of human serum specimens was reviewed and approved by the University of Texas Medical Branch (UTMB) Institutional Review Board (IRB number 20-0070). No informed consent was required since these deidentified sera were leftover specimens before being discarded. No diagnosis or treatment was involved either.

### Cells

Vero E6 (ATCC® CRL-1586) was purchased from the American Type Culture Collection (ATCC, Bethesda, MD), and maintained in a high-glucose Dulbecco’s modified Eagle’s medium (DMEM) containing 10% fetal bovine serum (FBS; HyClone Laboratories, South Logan, UT) and 1% penicillin/streptomycin at 37°C with 5% CO_2_. Culture media and antibiotics were purchased from ThermoFisher Scientific (Waltham, MA). The cell line was tested negative for mycoplasma.

### Human Serum

Three panels of human sera were used in the study. The first panel consisted of 25 pairs of sera collected from individuals 3-8 months after vaccine dose 3, and no more than 3 months after dose 4 of Pfizer or Moderna vaccine. This panel had been tested negative for SARS-CoV-2 nucleocapsid protein expression using Bio-Plex Pro Human IgG SARS-CoV-2 N/RBD/S1/S2 4-Plex Panel (Bio-rad). The second serum panel (n=29) was collected from individuals who had received 2 doses of mRNA vaccine and subsequently contracted Omicron BA.1. The third serum panel (n=38) was collected from individuals who had received 3 doses of mRNA vaccine and subsequently contracted Omicron BA.1. The genotype of infecting virus was verified by the molecular tests with FDA’s Emergency Use Authorization and Sanger sequencing. The de-identified human sera were heat-inactivated at 56°C for 30 min before the neutralization test. The serum information is presented in Table S1-3.

### Recombinant Omicron sublineage spike mNG SARS-CoV-2

Recombinant Omicron sublineage BA.1-, BA.2-, BA.2.12.1-, BA.3-, BA.4/5-spike mNG SARS-CoV-2s that was constructed by engineering the complete spike gene from the indicated variants into an infectious cDNA clone of mNG USA-WA1/2020 were reported previously^10,13^. BA.2.75-spike sequence was based on GISAID EPI_ISL_13521499. Figure 1A depicts the spike mutations from different Omicron sublineages. The full-length cDNA of viral genome bearing the variant spike was assembled via in vitro ligation and used as a template for in vitro transcription. The full-length viral RNA was then electroporated into Vero E6-TMPRSS2 cells. On day 3-4 post electroporation, the original P0 virus was harvested from the electroporated cells and propagated for another round on Vero E6 cells to produce the P1 virus. The infectious titer of the P1 virus was quantified by fluorescent focus assay on Vero E6 cells and sequenced for the complete spike gene to ensure no undesired mutations. The P1 virus was used for the neutralization test. The protocols for the mutagenesis of mNG SARS-CoV-2 and virus production were reported previously^13^.

### Fluorescent focus reduction neutralization test

A fluorescent focus reduction neutralization test (FFRNT) was performed to measure the neutralization titers of sera against USA-WA1/2020, BA.1-, BA.2-, BA.2.12.1-, BA.3-, and BA4/5-spike mNG SARS-CoV-2. The FFRNT protocol was reported previously.^9^ Vero E6 cells were seeded onto 96-well plates with 2.5×10^4^ cells per well (Greiner Bio-one™) and incubated overnight. On the next day, each serum was 2-fold serially diluted in a culture medium and mixed with 100-150 focus-forming units of mNG SARS-CoV-2. The final serum dilution ranged from 1:20 to 1:20,480. After incubation at 37°C for 1 h, the serum-virus mixtures were loaded onto the pre-seeded Vero E6 cell monolayer in 96-well plates. After 1 h infection, the inoculum was removed and 100 μl of overlay medium containing 0.8% methylcellulose was added to each well. After incubating the plates at 37°C for 16 h, raw images of mNG foci were acquired using Cytation™ 7 (BioTek) armed with 2.5× FL Zeiss objective with a wide field of view and processed using the software settings (GFP [469,525] threshold 4000, object selection size 50-1000 µm). The fluorescent mNG foci were counted in each well and normalized to the non-serum-treated controls to calculate the relative infectivities. The FFRNT_50_ value was defined as the minimal serum dilution to suppress >50% of fluorescent foci. The neutralization titer of each serum was determined in duplicate assays, and the geometric mean was taken. Tables S1-3 summarize the FFRNT_50_ results.

### Statistics

The nonparametric Wilcoxon matched-pairs signed rank test was used to analyze the statistical significance in Figures 1-2.

## Data availability

The raw data that support the findings of this study are shown in the Source data file.

## Acknowledgments

We thank colleagues at the University of Texas Medical Branch (UTMB) for helpful discussions. We thank Michael L O’Rourke from the Information System Department at UTMB for assisting with electronic medical record systems. P.-Y.S. was supported by NIH grants HHSN272201600013C and U01AI151801, and awards from the Sealy & Smith Foundation, the Kleberg Foundation, the John S. Dunn Foundation, the Amon G. Carter Foundation, the Gilson Longenbaugh Foundation, and the Summerfield Robert Foundation.

## Author contributions

Conceptualization, X.X., P.R., P.-Y.S.; Methodology, X.X., J.Z., C.K., M.L., P.R., P.-Y.S; Investigation, X.X., J.Z., C.K., M.L., P.R., P.-Y.S; Resources, X.X., P.R., P.-Y.S; Data Curation, X.X., J.Z., C.K., M.L., P.R.; Writing-Original Draft, X.X., J.Z., R.R., P.-Y.S; Writing-Review & Editing, X.X., P.R., P.-Y.S.; Supervision, X.X., P.R., P.-Y.S.; Funding Acquisition, P.-Y.S.

